# Shaping the zebrafish myotome by differential friction and active stress

**DOI:** 10.1101/505123

**Authors:** S. Tlili, J. Yin, J.-F. Rupprecht, G. Weissbart, J. Prost, T. E. Saunders

## Abstract

Organ formation is an inherently biophysical process, requiring large-scale tissue deformations. Yet, understanding how complex organ shape emerges during development remains a major challenge. During fish embryogenesis, large muscle segments, called myotomes, acquire a characteristic chevron morphology, which is believed to play a role in swimming. The final myotome shape can be altered by perturbing muscle cell differentiation or by altering the interaction between myotomes and surrounding tissues during morphogenesis. To disentangle the mechanisms contributing to shape formation of the myotome, we combine single-cell resolution live imaging with quantitative image analysis and theoretical modeling. We find that, soon after its segmentation from the presomitic mesoderm, the future myotome spreads across the underlying tissues. The mechanical coupling between the myotome and the surrounding tissues is spatially varying, resulting in spatially heterogeneous friction. Using a vertex model, we show that the interplay of differential spreading and friction is sufficient to drive the initial phase of myotome shape formation. However, we find that active stresses, generated during muscle cell differentiation, are necessary to reach the acute angle of the myotome observed in wildtype embryos. A final ingredient for formation and maintenance of the chevron shape is tissue plasticity, which is mediated by orientated cellular rearrangements. Our work sheds a new light on how a spatio-temporal sequence of local cellular events can have a non-local and irreversible mechanical impact at the tissue scale, leading to robust organ shaping.

---

The formation of complex organ shape requires the integration of genetic information [1–4] with mechanical processes such as directed cell division and rearrangements [5–11] and interactions between tissues [12]. The highly robust form of organs [7] suggests that forming a precise shape is essential. However, it remains an open question how different biophysical and genetic processes dynamically interact during organogenesis [13].

In the zebrafish embryo, precursors of myotomes, somites, start to bend into a chevron shape soon after segmentation [14]. Posterior trunk and tail somites emerge from the presomitic mesoderm (PSM), Fig. 1**a**., whilst anterior counterparts are generated from the mesoderm during gastrulation. Somites are specified by periodic segmentation around every 30 min [14–16] and they give rise to slow and fast-twitch muscle fibres, dermomyotome and various types of progenitor cells [17–21]. The developmental stage of a specific somite is denoted by SN, where N counts the number of already formed somites, with a newly specified somite denoted as stage S1, Fig. 1**a**. The final myotome consists of slow muscle fibres, whose progenitors are initially located close to the notochord, and multinucleated fast fibres, whose progenitors are initially located more laterally, Fig. 1**b**.

**Figure 1.**
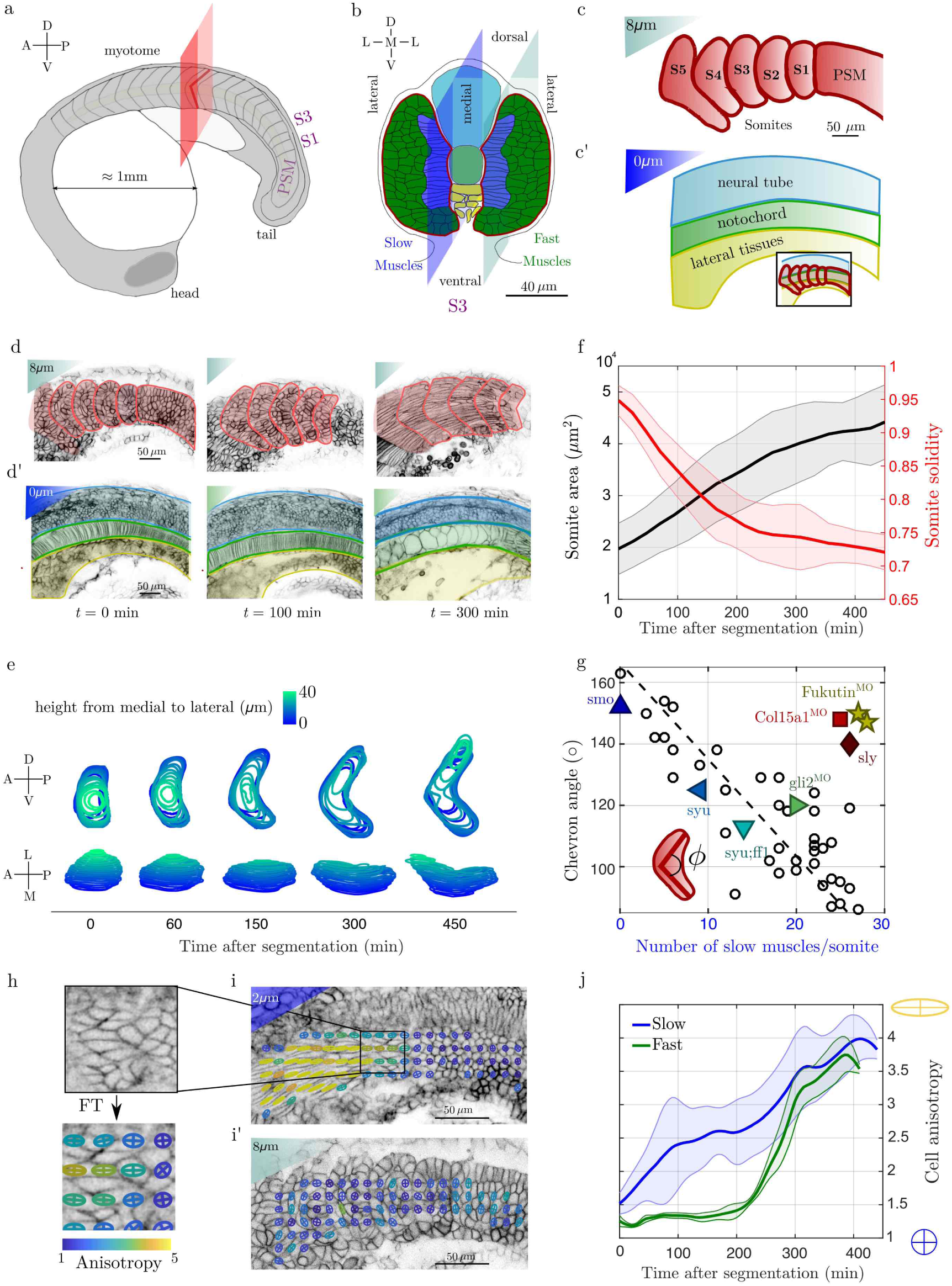
The chevron architecture of the myotome emerges early after segmentation from the PSM. (**a**) Sketch of a 21-somite zebrafish embryo. Red plane: transverse plane to the anterior-posterior direction. Two somites, at stages S1 and S3, are highlighted. (**b**) Sketch of a S3-stage somite (*t* ≈ 90 min post segmentation) in the transverse plane; dark blue (light green) cells are future slow (fast) muscle cells respectively. The dark and light blue planes represent the cross-sectional views shown in c and d. The notochord is at the centre (grey circle), with the neural tube located more dorsally. Ventral tissues not shown for clarity. (**c-c’**) Cartoon of somite shape in transverse view (c, plane lying *z* = 8*μ*m from notochord) and underlying tissues (**c’,** plane crossing the notochord, neural tube and ventral tissues. Inset shows shape of somites superimposed on underlying tissues. (**d-d’**) Confocal images and superimposed contours of (d) somites and PSM (red lines) and (**d’**) neural, notochord and mesoderm tissues (blue, green and yellow lines, respectively) at *t* = 0,100, 300 min post segmentation from PSM for the central somite shown in first panel. (**e**) 3D evolution of somite shape after segmentation from PSM of a representative wildtype embryo shows spreading of somite in DV-axis and emergence of chevron shape at ~ 150 min. (**f**) Cross-sectional area and solidity (i.e. the ratio of the somite area over the area of its convex hull) of segmented somites for the most medial layer of future fast muscle fibres (as in d) as a function of time after segmentation. Shaded regions represents ±1*s*.*d*.. (**g**) Chevron angle (in degrees) in the layer of most medial future fast muscle fibres against number of slow muscle cells per chevron. Black circles: Cyclopamine treated embryos at different concentrations. Triangles: morpholinos and mutants affecting cell differentiation (dark blue up Δ: smo [57], light blue left ⊲: syu [58], cyan down V: syu+ff1 [58], and green ⊳: gli2^*M0*^ [38]). Morpholinos or mutants altering tissue integrity (dark yellow star * : Fukutin [34], light red square □: Col15a1a^*MO*^ [33], dark red diamond ◊: sly [29, 59]). See Methods Table 1 for further details. (**h**) Fourier transform image analysis method provides a cell elongation field, with the anisotropy represented by ellipsoids (Methods). Cell elongation is along the major axis of the ellipse. (i) Elongation map of future slow (**i**) and fast (**i’**) muscle fibres. (**j**) Mean cell anisotropy as a function of time post segmentation for future slow muscle fibres (i, blue) and for the layer of most medial future fast muscle fibres (**i’,** green). Shaded regions represent ±1*s.d*.. In (**f**) and (**j**): average is performed over 11 somites from 6 embryos.

The mature myotome has a distinctive V (“chevron”) shape, [14], Fig. 1**a**, which is thought to be important for swimming [22]. A number of hypotheses have been proposed to explain chevron formation, including roles for: the swimming motion itself [23]; older myotome segments acting as templates for younger segments [24, 25]; tissue shear flow between the notochord and the developing myotome [26]; the interplay between intra-segmental tension and fixed myotome boundaries [14].

Here, we combine quantitative analysis of *in vivo* imaging data with modeling to show that a robust chevron shape emerges from the interplay between short-ranged processes (including cell differentiation and cell neighbour exchanges) and long-ranged mechanical processes mediated by the coupling between developing somites and their surrounding tissues.

## Symmetry breaking in the somite occurs early after segmentation from the PSM

We imaged somites at subcellular resolution inside the developing embryo, from their segmentation within the PSM (earliest S-2) to mature myotome stage (S5 onward), Fig. 1**c-d**, Methods, Supplementary Movie 1. Immediately after segmentation, somites are approximately cuboidal [27, 28], Fig. 1**d-e** and Supplementary Movie 2. Quantifying the somite contours over 8 h, we observe that the process of chevron formation occurs during phases S1 to S5, Fig. 1**e** and Supplementary Fig. 1. Somite volume is approximately constant during the 7 h following segmentation (Supplementary Fig. 2c). Immediately after segmentation the somites begin to change shape, with flattening in the medial-lateral (ML) axis, leading to an increased contact area with the medially underlying tissues (notochord, neural tube and ventral tissues), Fig. 1**e-f**, Supplementary Movie 2 and Supplementary Fig. 1.

Concurrently with spreading, we observe symmetry-breaking in the somite shape along the anterior-posterior (AP), Fig. 1**e**. In the medial region a “U” shape emerges that always points toward the anterior of the embryo. This “U” subsequently sharpens into the chevron shape, Fig. 1**e**.

## Chevron angle is impacted by both internal and external factors to the myotome

The shape of the myotome is known to be sensitive to a range of perturbations [29, 30], including to: (i) signaling pathways [31, 32]; (ii) the surrounding extracellular matrix (ECM) [33, 34]; and (iii) the surrounding tissues [35, 36]. Under perturbation, the myotome becomes more “U”-like or even flat. We are unaware of perturbations that sharpen the chevron, suggesting that the shape is tightly controlled and may be evolutionarily optimised. We quantified the chevron angle in a range of different conditions, using genetic (*smo*^-/-^, Supplementary Movie 3) and drug perturbations (Cyclopamine, Shh pathway inhibitor, Supplementary Movie 4). We complemented this with data from the literature, Fig. 1**g** and Methods Table 1. The chevron angle increases linearly from 90^*o*^ towards 180^*o*^ with decreasing slow muscle number. In contrast, altering of the extracellular matrix at the interfaces of somites and axial tissues (*e.g*. through Col15a1a^*MO*^, Fukutin^*MO*^ or *lamcl*^-/-^, see Methods Table 1) drastically reduces the chevron angle while the number remains largely unchanged, Fig. 1**g**. These results suggest that both muscle cell differentiation (intrinsic to each somite) and ECM interactions (at the interface between somites and surrounding tissues) are critical in forming the chevron.

## Somite deformation occurs prior to fast muscle fibre elongation

Concurrent with the tissue shape changes, cells within the somite begin differentiation into specific muscle fibres [17, 20, 21, 27, 37, 38]. The most-medial layer of cells undergoes differentiation into slow muscle fibres at the onset of somite segmentation, Fig. 1**b** [21]. Slow muscle fibres, which are epithelial-like before segmentation, rapidly elongate along the AP-axis until they span the somite compartment. To quantify the dynamics of slow muscle elongation, we used a Fourier transform method to analyse the evolution in cellular anisotropy within the somite, Fig. 1**h-i**, Methods [39]. We find that signatures of future slow muscle fibre elongation are apparent even before segmentation, and that these cells rapidly extend over the next 100 min, Fig. 1**j**. In contrast, fast fibres elongation occurs significantly later, at around 250 min, Fig. 1**j**. Comparing with Fig. 1**e**, we see that the chevron is apparent around 200 min after segmentation, yet fast fibres only fully elongate around this time. Despite fast muscle fibres representing > g80% of somitic cells, the future myotome acquires the characteristic chevron shape before most of these cells have begun to elongate.

## Spatio-temporal variation in somite-tissue coupling correlates with the chevron shape

As perturbations to surrounding tissues and ECM alter the myotome shape, we explored the mechanical coupling between somites and surrounding tissues. We used 2D optic flow to quantify the cellular velocity fields inside the somites (at different medial-lateral locations) and in the adjacent notochord and neural tube, Fig. 2**a-b**, Methods. We computed the averaged in-plane 2D velocity fields in the medial-lateral (ML) axis, Supplementary Fig. 3, as the shear velocities along the ML axis were comparatively small.

**Figure 2.**
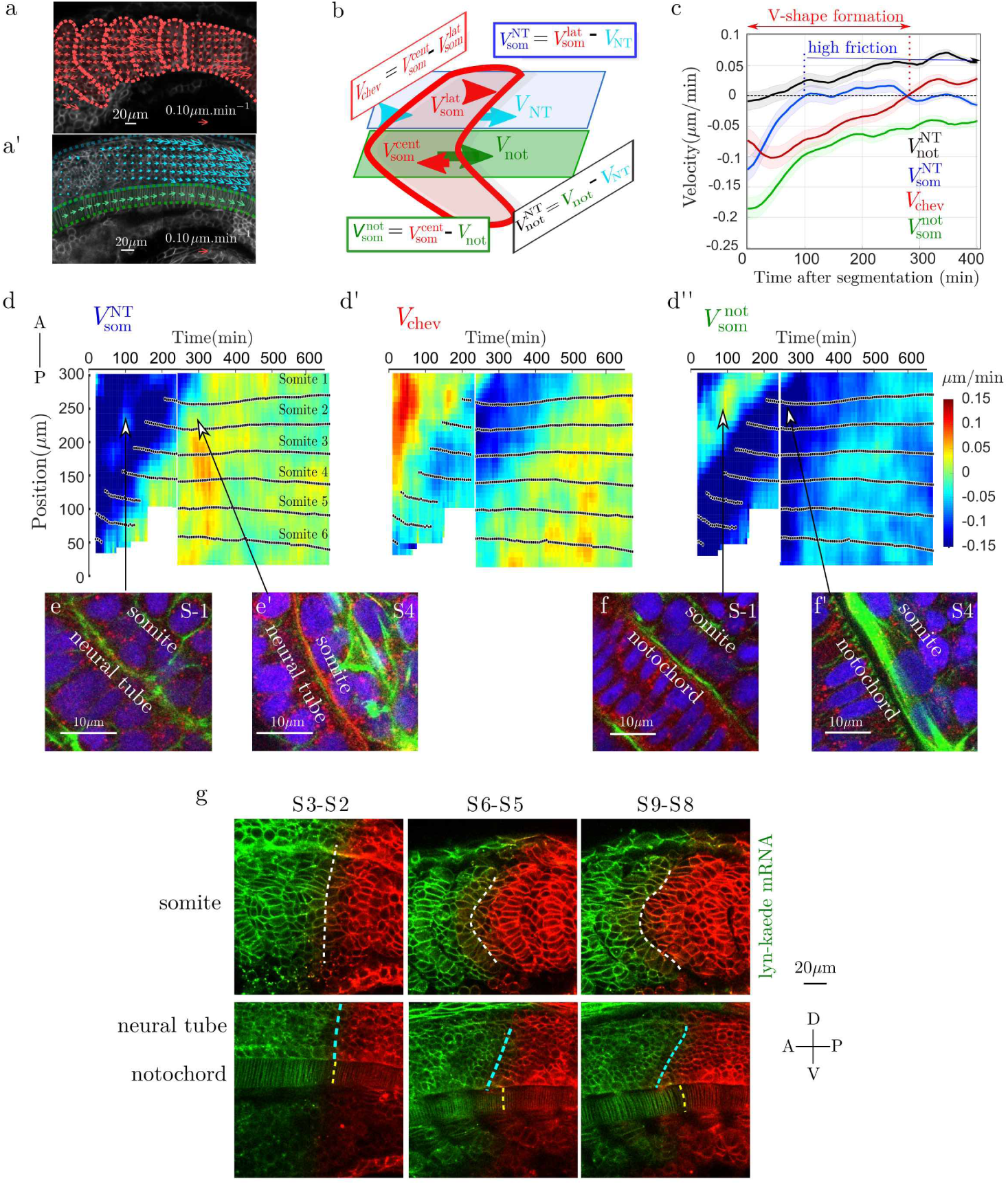
Differential tissue flow and heterogeneous mechanical coupling between tissues correlates with the emergence of a chevron shape. (**a-a’**) Velocity fields estimated by Optic Flow (Methods) within the somite (red arrows, a), neural tube (cyan, **a’**) and notochord (green, **b’**). (**b**) Definition of the mean anterio-posterior (AP) velocities within each tissues: neural tube *(NT,* cyan), notochord *(not,* green) and somites *(som,* red). (c) Evolution of the relative tissue AP velocities (average performed on *n* = 11 somites from *N* = 5 embryos) after segmentation from the PSM. Negative values of the shear strain rate *V*_chev_ represent the period of chevron shape emergence. Near zero values of 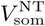 for *t >* 100 min post-segmentation indicate the onset of a large friction between the notochord and somites. Shaded regions represent ±1*s*.*d*. (**d-d”**) Kymographs of shear velocities 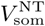, *V*_chev_ and 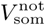 shows somite-to somite reproducibility of the features identified in (c). Each panel from two embryos, with stitching at *t* = 220 min. Black dots indicate the position of each somite centre of mass along the AP axis, with somite labelling representing somite number with respect to the start of the movie. In (**d’**), negative shear (blue coloured region) indicates the region where the chevron shape emerges in the somite. (**e-f**) Confocal images of actin (green), laminin (red) and nuclei (blue) in the transverse plane to the AP-axis for somites *S*-1 and *S*4 (scale bar: 10 *μ*m); (**e**) Closeup of the somite/neural tube interface. Arrows highlight correlation between actin and laminin localisation with the corresponding tissue-tissue flows shown in **d**. (**f**) Closeup of the somite/notochord interface. Arrows highlight correlation between actin and laminin localisation with the corresponding tissue-tissue flows shown in **d”.** (**g**) Lyn-Kaede showing relative movement of the somites (top) with respect to the underlying notochord and neural tube (lower) from S2 to S9, with photo-switching of Kaede performed at S2 stage in the more posterior somites, Methods. Dashed lines highlight interfaces between the two fluorescent regions.

To gain insight into the physical coupling between tissues, we focused on relative tissue velocities. We primarily considered the velocity component along the AP-axis for each tissue (*V*_not_ (notochord), *V*_som_ (somite) and *V*_NT_ (neural tube)), Methods. We define the shear velocity within the somite *V*_chev_ as the relative difference in the velocity of cells at the DV-midline and of those in more dorsal positions, Fig. 2**b**, along with similar shear velocities between somites and surrounding tissues: 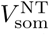 (relative somite velocity compared to neural tube); and 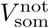 (relative somite velocity compared to notochord). Lastly, we define a shear velocity 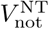 between the notochord and neural tube, Fig. 2**b**.

Each of these shear velocities has distinct behaviour, Fig. 2**c**, kymographs Fig.**2d-d”** and Supplementary Fig. 4. In agreement with the chevron formation timescale identified in Fig. 1**f**, *V*_chev_ < 0 during the first 5 h after segmentation, Fig. 2**d**’. Within this time, the notochord moves more posteriorly than the neural tube during chevron formation, as 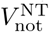 remains positive after segmentation, Fig. 2**c**. Hence, somites are not passively deformed by an underlying tissue shear, in which case the chevron would point toward the embryo posterior. Soon after segmentation 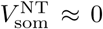, implying that the somite and neural tube move concomitantly, Fig. **2c,d**. Similarly, before segmentation, future somites and notochord move concomitantly, Fig.**2d”**. In contrast,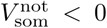 throughout the 6 h after segmentation, Fig.**2c,d”,** implying that somites move in the anterior direction relative to the notochord.

We complemented this analysis with live imaging of embryos injected with lyn-Kaede, Methods. By switching the Kaede at a somite boundary, we observed the differential movement of the notochord and neural tube with respect to the somite, Fig. 2**g**, consistent with the above velocity maps. From these observations, our hypotheses are: (i) that somites and neural tube are weakly mechanically coupled prior to segmentation but strongly linked after segmentation; (ii) that somites and notochord are mechanically coupled prior to segmentation and uncoupled afterwards.

## Mechanical coupling between tissues varies in time

To explore whether temporal changes in the relative movements between tissues are correlated with changes in physical interactions between these tissues, we examined the distribution of actin and the ECM component laminin, [40], Fig. 2**e-f** and Supplementary Movie 5-6. Before segmentation, future somitic cells are in contact with the notochord, Fig. 2**f**, while after segmentation, a gap between these cells and the notochord emerges, together with the appearance of large actin fibres. Such loss of contact suggests a reduced friction between the notochord and the slow muscle fibres.

In contrast, cells in the PSM appear in contact with the neural tube, with progressive actin accumulation between the tissues, Fig. 2**e**. After segmentation, a layer of laminin appears between the somitic cells and the neural tube, Fig.**2e’**, which suggests that mechanical coupling between the neural tube and somitic cells further increases. Other molecules could also contribute to adhesion, *e.g.* integrin and fibronectin, whose localisation are tightly regulated during somite formation [41].

Following [42, 43], we expect strongly (weakly) adhered tissues to have a high (low) effective interfacial friction coefficient. Such a framework has proved fruitful in understanding tissue-tissue interactions during early zebrafish morphogenesis [44]. Below, we incorporate this idea -along with somite spreading and cell differentiation-within a vertex model to test how tissue-tissue coupling drives the chevron shape of the myotome.

## Simulating tissue shape formation within a vertex model

The chevron first emerges on the medial side of the somite, which includes slow muscle fibres and the most-medial future fast fibres. We simulate an average 2D layer of cells located within this medial compartment of each somite, Fig 3**a**. We do not distinguish specific muscle types.

**Figure 3.**
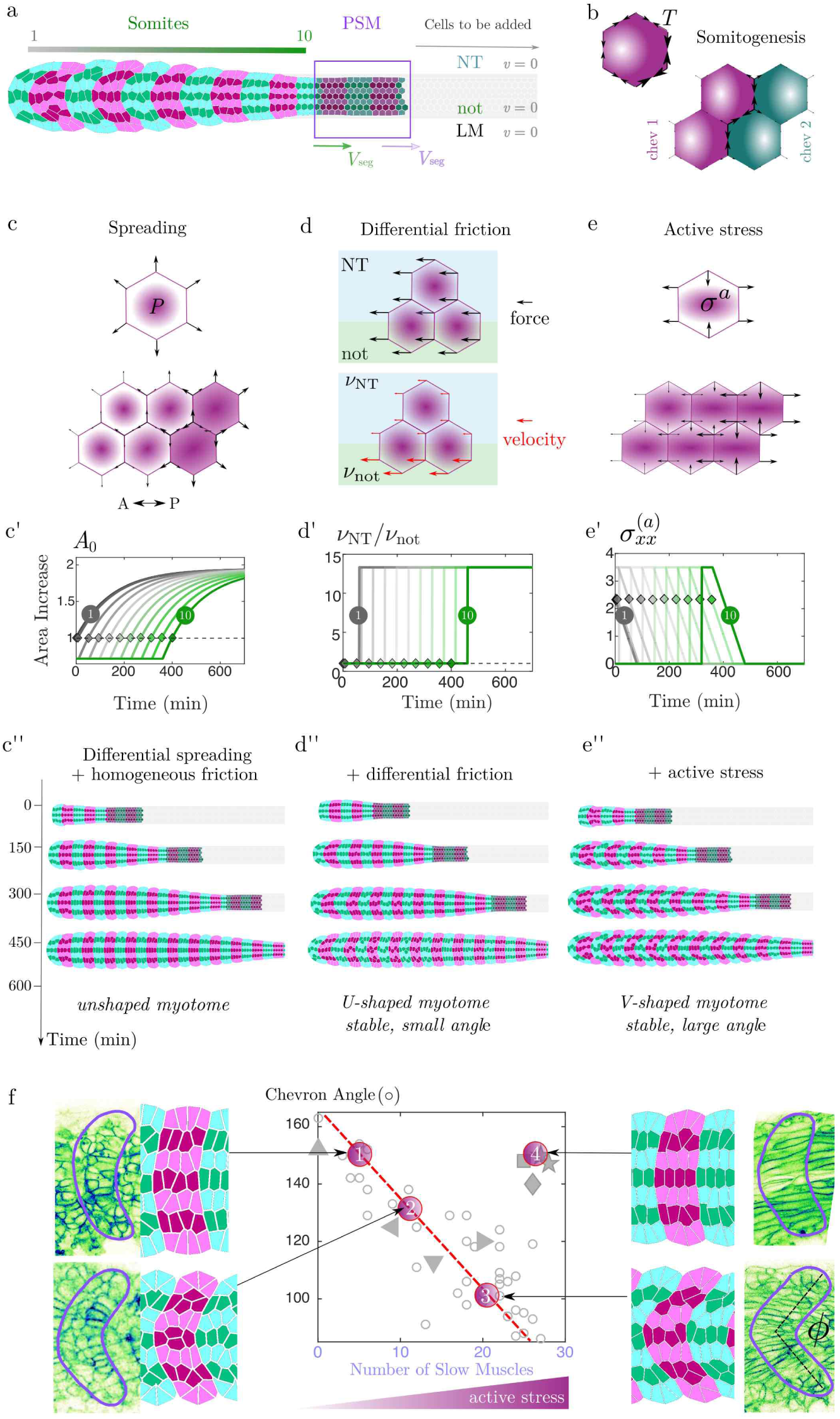
The chevron shape emerges in a vertex model incorporating somite spreading, differential friction and active stress. (**a**) Simulated geometry used in model. The number of simulated cells increases with time as new cells are progressively added from the tailbud region (highlighted by purple region); magenta (green) cells belong to somite number 2N (*2N* + 1) respectively. New somites appear at a velocity *V*_seg_ = 1 somite / (30 min). (**b-e**) Principle elements included within the vertex model: (**b**) Somite segmentation is implemented through an increased tension at the somite compartment boundaries. (c) Differential spreading is implemented through a wave of increased cellular pressure along the AP axis, leading to a spatial modulation of outward forces (black arrows). (**c’**) Exponential increase in the somite target area as a function of time, based on experimental measurements. Grey curve (dark green) corresponds to first (last) somite formed in simulation (diamonds indicate timing of segmentation of specific somite from PSM, **a**). (**c”**) Simulations with differential spreading only (*i.e.* homogeneous friction): somites do not buckle. (**d**) The vertex displacement (red arrow) is spatially modulated by an inhomogeneous friction coefficient *v*, where *v* = *v*_NT_ = *v*_LM_ for vertices over the neural tube and ventral tissues; and *v* = *v*_not_ otherwise. (**d’**) The ratio of friction between the somite and neural tube and the somite and notochord, implemented as a step function (related to Fig. 3**e-f**). (**d”**) Simulations with somite spreading and differential friction: somites fail to form a long-ranged sharp chevron shape. (e) An imposed bulk active stress *σ*^(*a*)^ leads to elongation forces along the AP axis (black arrows). (**e’**) Active stress is set to be maximal for each somite soon after segmentation, corresponding to slow muscle fibre elongation. (**e”**) Simulations with active stress and differential friction (wildtype case): somites acquire a stable chevron shape. (**f**) Comparison of experimentally measured chevron angle (Fig. 1**g**) with the angle measured for four simulation outcomes. Only the active stress level is varied from points 1 to 3 (all other parameters fixed), describing embryos treated with 50 *μ*mol (1), 10 *μ*mol (2) of Cyclopamine and wildtype embryos (3). (4) corresponds to the homogeneous friction case, describing the perturbed tissue-tissue coupling of Col15a1a^*MO*^.

Each cell is described by a polygon whose summits, called vertices and denoted by *X_i_*, correspond to the edges of cell-cell interfaces. Cellular movements and deformations are described through the dynamics of the cell vertices, which is set by the following force-balance equation:

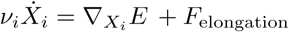

This equation has three critical elements. (i) spatially-dependent friction: *v_i_* represents the friction on vertex i exerted by the underlying tissues, (ii) active stress forces, denoted *F*_elongation_ which are generated by the elongation of slow muscle fibres, Methods, and (iii) cell-scale forces regulating cell shape, ∇_*Xi*_*E*. Following [45], we consider

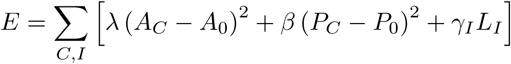

where *A*_0_(*P*_0_) represents the preferred area (perimeter) of a cell *C,A_C_*(*P_C_*) the actual area (perimeter) of a cell at a given time, and *L_I_* the length of cell-cell interface *I*, Methods. λ represents the pressure forces involved in cell area regulation, while *β* and γ*I* represent the strength of cell- and interface-dependent tensions respectively. Following [46], we introduce stress fluctuations through stochastic modulation in the tension of each cell-cell contact, Methods. After segmentation, no cellular exchanges with neighbouring somites are observed.

We model this compartmentalisation by increasing the tension γ along the somite/somite boundaries, Fig. 3**b**, Methods. Lastly, to simulate growth and division within the PSM and tailbud, we continuously add new cells at the posterior end of the tissue, at a rate determined by the segmentation clock [16].

## Somite spreading and differential friction are sufficient to generate a shallow chevron shape

We first tested in the model the effects of spatially varying friction and somite spreading. To simulate the wave of spreading we varied the target area of each cell 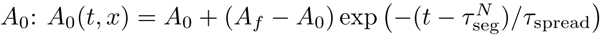 with *τ*_spread_ = 200 min extracted from experiment, Fig. 3**c-c**’, and 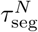 is the segmentation date of the *N*-th somite. During spreading, each cell has a constantly increasing target area and hence exerts pushing forces on its neighbouring cells. We first considered somite spreading with uniform friction, Fig. 3**c**”. Along the DV axis of the somite, all cells have the same target area and spread together. However, along the AP axis the cells are not at the same stage of spreading. Newer (and subsequently smaller) somites have a higher spreading rate than more anterior (older) somites, resulting in a net force along the central part of more anterior somites and a slight bending towards the head occurs. However this bending is insufficient to irreversibly deform the somites; they relax once spreading is finished, Supplementary Movie 7.

We next introduced spatially inhomogeneous friction within the model, Fig.**3d-d’**; *v_i_* depends on vertex position *X_i_.* After segmentation, we increase the friction coefficient over the neural tube and ventral tissues while decreasing the friction coefficient over the notochord, Fig. 3**d’**. Combining spreading with non-uniform friction gives rise to clear symmetry breaking, with somites deforming into a shallow chevron, Fig. 3**d”** and Supplementary Movie 8. As cells lying above the notochord slide faster than those located more dorsally, the stress associated to the somite spreading creates a DV-shear that deforms somites into a U shape. Such tissue deformation also alters individual cell shapes. If the tissue is sufficiently plastic (*i.e.* frequent cell rearrangements), then cell rearrangements relieve stress induced by the shape changes, resulting in a sharpening of the somite boundary and the emergence of a stable but shallow chevron in early somites. However, this shape does not propagate to younger somites as tissue spreading is insufficiently rapid to trigger cell rearrangements in later segments. Therefore, incorporating realistic parameters (derived where possible from experiments) within such a model cannot generate a sharp chevron similar to wildtype embryos.

## Active stress due to muscle fibre differentiation modulates chevron angle

Slow muscle fibres start to elongate soon after somite segmentation from the PSM, Fig. 1**j**. Such elongation likely exerts a shear stress on the more lateral layers of future fast muscle fibres. To model the mechanical constraints imposed by the layer of slow muscle fibre, we used active gel theory, which is a hydrodynamic description of the acto-myosin cortex that encompasses contractility and filament polymerisation through a local active stress tensor [47], Methods. We considered a traceless active stress to discriminate its contribution from somite spreading. The positive component (extension) of the active stress is orientated along the AP-axis, in line with muscle fibre elongation, with a corresponding negative component (contraction) orientated along the DV-axis, Fig. 3**e**. We assume that the active stress is maximal at the start of slow muscle elongation, with a further linear decrease to zero by the end of slow muscle elongation, Fig. 3**e**’, which leads to a convergence (DV-axis)-extension (AP-axis) wave within the somites. The inclusion of such orientated active stress is then sufficient to shape the tissue into a stable and sharp chevron shape, Fig. 3**e**” and Supplementary Movie 9.

## Model predictions for myotome shape under perurbations

As shown in Fig. 1**g**, the chevron shape changes with slow muscle number. Within our model, this corresponds to changing the active stress, but leaving other components unchanged. Without active stress, the model predicts a transient shape deformation in the somite before relaxing. These dynamics are strikingly similar to smo^-/-^ embryos, where there is no slow muscle specification, Supplementary Movie 3. Intermediate levels of active stress in the model result in reduced chevron angle, as observed experimentally, Fig. 3**f**. Perturbations to the interactions between tissues (*e.g*. Col15a1a^*MO*^) likely change tissue-tissue coupling, thereby reducing the effects of differential friction. Reducing friction heterogeneity within the model (while keeping active stress) results in a mild but stable bending of the myotome, consistent with experiments, Fig. 3**f** and Supplementary Movie 10.

### Dynamics of chevron formation

We next challenge the model capacity to reproduce the tissue dynamics observed during chevron formation. In both experiments and simulations, we quantified the tissue velocity, Fig. 4**a**, anisotropic strain rate (ASR), Fig. 4**b** and isotropic strain rate, Supplementary Fig. 6. By definition, the ASR provides the local tissue expansion direction. Common features between the ASR fields in experiments and simulations (Fig.**4a’**,**b’**) are: (i) in S1, somites undergo DV-convergence and AP-extension (purple bars, Fig. 4**b**), correlating with the onset of slow muscle elongation, Fig. 1**j**; (ii) in S2 somites, the ASR is near zero, (iii) from S3 to S6, somites undergo shear between central and lateral regions.

**Figure 4.**
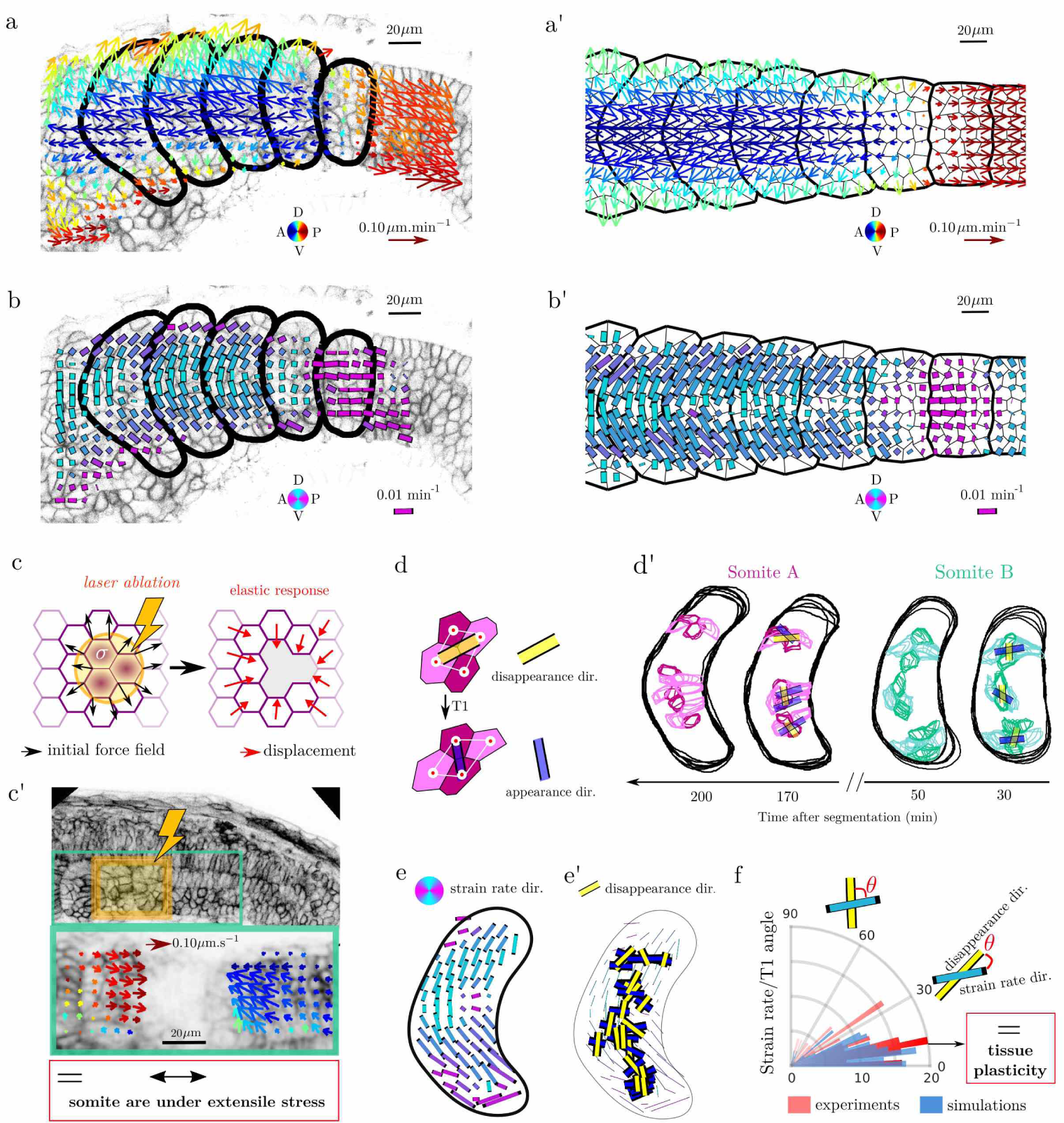
Model accurately predicts forces within the somite and orientation of cellular rearrangements. (**a-a’**) Comparison of velocity field in (**a**) experiments and (**a’**) simulations, measured using optic flow, Methods. Arrow colour represents direction and length represents speed. (**b-b’**) Comparison of the anisotropic component of the strain rates (ASR) in (**b**) experiments and (**b’**) simulations (magenta: AP orientation; cyan: DV orientation). Bar color represents orientation and length represents the strain rate magnitude (see Methods). (**c**) Cartoon of predicted relaxation direction upon ablation of somitic tissue. (**c’**) (Top) Laser ablation (yellow box) of somites at stage S0 and S1. (Bottom) Zoomed in region highlighted above, with arrows representing tissue velocity from optic flow analysis in the 10 s after ablation. Colour coding as a). (**d**) Scheme of cellular rearrangements, with cells losing contact joined by yellow bar and cells forming new contacts by blue bar. (**d’**) Experimental examples of 3D cellular rearrangements at different somite stages for 2 somites. (**e**) Time and ensemble averaged ASR (*n* = 4 somites). (**e’**) Accumulated cell rearrangements orientations (across 4 somites) superimposed on the ASR map. (**f**) Rose plot alignment of cellular rearrangement with ASR in experiments (*n* = 44 from 4 somites) and simulations (*n* = 60 from 6 simulated somites).

Our interpretation of the ASR field is that active elongation of slow muscle cells is maximal at somite S1, generating an extensile stress along the AP-axis that compresses both the PSM and anterior somites. Such compression pattern is similar to the one produced by a single cell actively extending in a passive tissue, Supplementary Fig. 5, yet at the larger tissue level. To test this interpretation, we laser ablated a region of newly formed somitic tissue, Fig. 4**c’,** Supplementary Movie 11 and Methods. We observed a rapid relaxation of neighbouring tissues towards the ablated tissue, confirming that newly segmented somites are pushing their neighbours.

We then investigated the role of tissue plasticity. In a purely elastic material, the somite shape would eventually relax, since the shear stresses generated by cell elongation and spreading are transient. A plastic/viscous-like behaviour is therefore required to acquire a stable chevron shape. Within the vertex model, we implemented passive cell rearrangements (Fig. 4**d**). Due to the shear forces emerging in the model, passive cellular rearrangements are naturally oriented along the ASR (Fig. 4**f**) indicating that the bulk somitic tissue has a plastic-like behaviour [48].

Experimentally, we observe that tissue flows do not generate large cell deformations, Supplementary Fig. 8, which suggests the existence of cell rearrangements [49, 50]. Cell divisions can also relax cell shape [49]; however, we found only infrequent cell divisions during myotome formation, with less than 10 % of cells dividing during the whole process.

We explicitly show how cell rearrangements occur by tracking cellular shapes in 3D inside S1, S2 and S3 somites using high temporal resolution movies, Fig. 4**d**’ and Supplementary Movie 12. To correlate them with the ASR, we superimposed the rearrangements in time over an ASR field map, Fig.**4e,e’** and Methods. Cellular rearrangements are indeed closely aligned with the ASR, Fig. 4**f**, in agreement with our theoretical predictions.

While intra-somite cell rearrangements are needed, inter-somite cell exchanges ought to be prevented to preserve the somite shape. Based on our simulations, we expect somite-somite interfaces to be rough in *tbx6*^-/-^ embryos, in which somites compartmentalisation is abolished. By using lyn-Kaede to define boundaries within *tbx6*^-/-^ embryos, we indeed see greater inter-compartmental mixing, Supplementary Fig. 7b-c. We note that it has been previously shown that using a heat-shock inducible Tbx6, somite shape can be rescued in *tbx6*^-/-^ embryos, showing that the chevron formation is an emergent property, i.e. that it is not due to a template mechanism [51].

## Conclusion

During myotome formation, somites are under mechanical stress from both internal (somite spreading, cell elongation) and external (tissue-tissue coupling) processes. Combining our experimental and cell-based numerical approaches, we propose the following sequence of mechanical events leading to chevron-shaped myotomes: (i) increased line tension between developing somites leads to mechanically segmented cell compartments, Fig. 3**b**; (ii) somite differential spreading (Fig. 3**c**) leads to a pressure gradient along the AP axis, which, combined with the onset of a differential friction along the DV axis (Fig. 3**d**), leads to a buckling instability; and (iii) muscle fibre elongation further contributes to buckling (Fig. 3**e**), which trigger cell rearrangements that maintain a stable chevron shape. Our 2D model incorporates features resulting from the 3D dynamics of the somite, yet neglects cell heterogeneities within the fast-muscle cell population, Supplementary Fig. 8. Though we cannot discount other possible mechanisms, our model is minimal yet sufficient to recapitulate the dynamics of somite shape formation in both wildtype and perturbed embryos.

Recent works have shown (i) how active stress can generate complex flows within *in vitro* tissues [53–55], (ii) how tissue-tissue friction affects tissue flows during early zebrafish embryogenesis [44] and (iii) how rheological properties set the shape of the zebrafish PSM and tailbud [46]. Here, we integrate these approaches in the vertex-model framework to understand the shaping of a functional organ in terms of the interplay between (i) active stresses generated by muscle cell differentiation, (ii) spatially heterogeneous friction and (iii) tissue plasticity.

It is interesting to compare with somite formation in other vertebrate systems, such as the chicken and mouse embryos whereby somites do not acquire a chevron shape. While in our model the notochord needs to be centred to generate the chevron shape, in mouse and chick embryos the notochord is off-centred and located towards the ventral border (see Supplementary Fig. 9). Given such tissue arrangement, we do not expect somites to buckle, even in the presence of differential tissue-tissue frictions [56]. Therefore, our work suggests that both tissue-tissue interactions and tissue positioning can play a key role in shaping organs.

## Acknowledgements

We thank Philip Ingham for support with zebrafish experiments. We acknowledge funding from a Singapore National Research Foundation Fellowship awarded to T.E.S. (grant No. 2012NRF-NRFF001-094), an HFSP Young Investigator Grant awarded to T.E.S. (grant no. RGY0083/2016), and the Mechanobiology Institute. Fish facilities were provided by the Institute of Molecular and Cellular Biology, Singapore. We thank MBI Science Communications for assistance with graphics as well as Olivier Hamant and Francois Graner for critical reading of the manuscript.

## Author contribution

J.Y. and T.E.S. planned the experiments, S.T. and T.E.S. planned the image analysis, and S.T., J.-F.R., J.P. and T.E.S. planned the theoretical analysis. J.Y. performed all experiments except laser ablation (by S.T.). S.T. performed the image analysis. S. T. and J.-F.R. developed the vertex model with inputs from G. W. and J.P., particularly regarding modelling of active stress. S.T., J.-F.R. and T.E.S. wrote the manuscript, with all authors contributing to manuscript preparation. We have no competing interests.

